# Microbial community analysis of biofilters reveals a dominance of either comammox *Nitrospira* or archaea as ammonia oxidizers in freshwater aquaria

**DOI:** 10.1101/2021.11.24.468873

**Authors:** Michelle M. McKnight, Josh D. Neufeld

## Abstract

Nitrification by aquarium biofilters transforms toxic ammonia waste (NH_3_/NH_4_^+^) to less toxic nitrate (NO_3_^-^) via nitrite (NO_2_^-^). Ammonia oxidation is mediated by ammonia-oxidizing bacteria (AOB), ammonia-oxidizing archaea (AOA), and the recently discovered complete ammonia oxidizing (comammox) *Nitrospira*. Prior to the discovery of comammox *Nitrospira*, previous research revealed that AOA dominate among ammonia oxidizers in freshwater biofilters. Here, we characterized the composition of aquarium filter microbial communities and quantified the abundance of all three known groups of ammonia oxidizers. Aquarium biofilter and water samples were collected from representative freshwater and saltwater systems in Southwestern Ontario, Canada. Using extracted DNA, we performed 16S rRNA gene sequencing and quantitative PCR (qPCR) to assess community composition and quantify the abundance of *amoA* genes, respectively. Our results show that aquarium biofilter microbial communities were consistently represented by putative heterotrophs of the *Proteobacteria* and *Bacteroides* phyla, with distinct profiles associated with fresh versus saltwater biofilters. Among nitrifiers, comammox *Nitrospira amoA* genes were detected in all 38 freshwater aquarium biofilter samples and were the most abundant ammonia oxidizer in 30 of these samples, with the remaining biofilters dominated by AOA, based on *amoA* gene abundances. In saltwater biofilters, AOA or AOB were differentially abundant, with no comammox *Nitrospira* detected. These results demonstrate that comammox *Nitrospira* play an important role in biofilter nitrification that has been previously overlooked and such microcosms are useful for exploring the ecology of nitrification for future research.

## Introduction

Aquaria are closed systems that can accumulate waste by-products of resident aquatic organisms. Nitrogenous waste in the form of ammonia/ammonium (NH_3_/NH_4_^+^), excreted by fish or produced from the degradation of organic matter, can exceed concentrations that are toxic to aquatic life, especially in newly established aquaria. Concentrations of unionized ammonia (NH_3_) greater than 0.1 mg/L can lead to chronic stress and disease of aquarium fish if not removed through nitrification or water changes (1, 2). To maintain a healthy aquarium, biofilters with established microbial communities that can perform nitrification are used to convert ammonia into nitrate.

The first step of nitrification, ammonia oxidation, can be performed by either ammonia-oxidizing bacteria (AOB; e.g., *Nitrosomonas*) or the more recently discovered ammonia-oxidizing archaea (AOA; e.g., *Nitrosotenuis, Nitrosopumilus*) (3). Nitrite oxidation is subsequently performed by nitrite-oxidizing bacteria (NOB; e.g., *Nitrobacter, Nitrospira*). Most recently, specific *Nitrospira* spp. were found to catalyze complete ammonia oxidation (comammox), possessing genes for both ammonia and nitrite oxidation. Comammox *Nitrospira* were first discovered in a deep oil exploration well and a recirculating aquaculture system (RAS) (4, 5) and were earlier predicted to be associated with biofilm growth exposed to low ammonia concentrations (6). All comammox bacteria discovered to date belong to a single genus, *Nitrospira*, which was previously considered a genus that contained only strict NOB (7), and they are classified into two distinct clades: comammox clade A and clade B (4). Since their discovery, comammox *Nitrospira* have been identified in many natural and engineered environments, including wastewater and drinking water treatment plants, soil, lake sediments, and groundwater-fed rapid sand filters (8–13). Currently, our overall understanding of factors that govern environmental distributions and abundances of comammox *Nitrospira* are unclear. However, several studies show that comammox *Nitrospira* are more abundant in habitats with low ammonia concentrations (9, 12) and kinetic studies have revealed that the cultivated comammox representative *Nitrospira inopinata* has a high affinity for ammonia (K_m(app)_ = 63 ± 10 nM NH_3_), suggesting that they have a competitive advantage in oligotrophic environments (14).

Initial aquarium filter studies concluded that AOB were solely responsible for ammonia oxidation in these systems (15, 16). Similarly, NOB were identified as being responsible for completing nitrification in association with AOB, with *Nitrospira* spp. serving as strict nitrite oxidizers (17). However, the first cultivated AOA representative, *Nitrosopumilus maritimus* SCM1, was isolated from gravel collected from a marine tropical aquarium (18), and several studies in subsequent years revealed a dominance of AOA in marine aquaculture biofilters (19, 20). A subsequent survey of ammonia oxidizers within biofilters collected from hobbyist freshwater aquaria, using qPCR analysis of the ammonia monooxygenase A gene (*amoA*), revealed that AOA were numerically dominant in freshwater aquaria compared to AOB (21). This survey also found that the relative abundance of AOA was negatively correlated with total ammonia concentration. Another study identified a dominance of AOA in freshwater aquarium biofilters, observing long-term stability within the sampled filter material (22). To date, no study has sampled aquarium biofilters to reassess the abundance of nitrifying microorganisms with the additional consideration of comammox *Nitrospira*. Given the identification of comammox *Nitrospira* in aquaculture systems (23), and the favourable high surface area for biofilm growth and low ammonia environment predicted for optimal growth of comammox *Nitrospira*, it is worth exploring their abundances in relation to other nitrifiers in these systems.

Here we revisit the microorganisms involved in aquarium nitrification, with an emphasis on ammonia oxidation, and determine the abundance of AOA, AOB, and comammox *Nitrospira* present in representative freshwater and marine aquarium biofilters, using qPCR analysis of the *amoA* gene. We also explore the total microbial community composition using 16S rRNA gene sequencing and evaluate aquarium characteristics that may influence microbial community composition. Based on relatively low ammonia concentrations typically present in home aquaria, and our previous AOA-focused aquarium data, we hypothesized that both comammox *Nitrospira* and AOA would dominate freshwater aquarium biofilter nitrifier communities.

## Methods

### Sample collection

Aquarium filter and water samples were collected in 2019 from members of the Kitchener Waterloo Aquarium Society (KWAS), community members, and local pet stores; a total of 38 freshwater and 8 saltwater tanks were sampled (Table S1-S2). Small pieces of sponge, foam, or floss (as appropriate for each filter) were collected from biofilters using sterile scissors and forceps and then stored in Ziplock bags. During transport to the University of Waterloo, samples were kept on ice and subsequently stored at -70°C prior to DNA extraction. Aquarium water samples were collected alongside filter samples and kept on ice during transport. Prior to storage at -20°C, water samples were aliquoted into two 50-mL Falcon tubes, one of which was filtered through a 0.45-μm syringe filter. Participants also provided aquarium information including temperature, maintenance history, water source, and the number and species of fish and plants (Table S1-S2).

### DNA extraction

DNA was extracted from aquarium biofilter samples using the DNeasy PowerSoil Kit (Qiagen). Filter samples were thawed and a ∼2 cm^3^ portion was cut into small pieces and placed into the 2 mL bead-beating tube using sterile scissors and forceps. The extraction was performed following the manufacturer’s protocol, with the addition of a 10-min pre-incubation at 70°C prior to bead beating. Subsequent homogenization was performed using a FastPrep-24 bead beater (MP Biomedical, Santa Ana, CA) for 45 s at 5.5 m s^-1^. Extracted DNA was visualized on a 1% agarose gel using Gel Red nucleic acid stain and quantified using Qubit dsDNA HS Assay Kit (ThermoFisher Scientific) and the NanoDrop 2000 (ThermoFisher Scientific). Extracted DNA samples were stored at -20°C until further use.

### Quantitative PCR

Quantitative PCR (qPCR) was performed on filter DNA extracts to determine both 16S rRNA and *amoA* gene copies. The 16S rRNA genes associated with bacteria were quantified using the primer set 341F/518R (24) and Thaumarchaeota/Thermoproteota 16S rRNA genes were amplified with primers 771F/957R (25). The *amoA* genes were amplified using comaA-244F(a-f) and comaA-659R(a-f) to target clade A comammox *Nitrospira* (12), crenamoA-23F/crenamoA-616R to target AOA (26), and amoA1F/amoA2R to target AOB (27). All qPCR amplifications were run as technical duplicates using the CFX96 Real-Time PCR Detection System (Bio-Rad, Hercules, CA, USA). All 10-μL reaction volumes contained 1X SsoAdvanced Universal SYBR Green Supermix (Bio-Rad, Hercules, CA, USA), 5 μg of bovine serum albumin, primers, and 1-10 ng of template DNA. The amount of forward and reverse primer used per sample for amplification varied depending on the primer set. For the crenamoA-23F/crenamoA-616R and amoA1F/amoA2R primer sets, 4 pmoles of each primer were used, the 341F/518R primer set used 3 pmoles of each primer, 2 pmoles of each primer were used for the 771F/957R primer set, and 5 pmoles were used for the comaA-244F(a-f) and comaA-659R(a-f) primer pair. The PCR conditions for the 341F/518R and 771F/957R primer sets involved an initial 3 min denaturation at 98°C, then 35 cycles at 98°C for 30 s followed by a combined annealing and extension step at 55°C for 45 s. For clade A comammox *Nitrospira amoA* gene amplification, an initial denaturation for 3 min at 98°C was also used, followed by 35 cycles at 98°C, 45 sec at 52°C, and 1 min at 72°C. Both the AOA and AOB *amoA* gene amplification began with an initial denaturation step for 3 min at 98°C, followed by 35 cycles of 30 sec at 98°C, 30 sec at 55°C or 60°C for the AOA and AOB amplification respectively, and 1 min at 72°C. Following all qPCR amplifications, a melt curve was run from 65-95°C with 0.5°C interval increases, each lasting for 2 sec. Gel-purified PCR amplicons generated from *Ca*. N. aquarius, *N. europaea*, and pooled aquarium biofilter DNA templates were used as qPCR standards for AOA, AOB, and comammox *amoA* gene targets, respectively, whereas purified PCR amplicons generated from *Thermus thermophilus* and *Ca*. N. aquarius DNA were used as standards for bacterial and *Nitrososphaeria* 16S rRNA gene targets, respectively (Table S3-S4). Starting DNA copy numbers in standards used for qPCR were determined based on the DNA concentrations of the PCR amplicons, with standard curves ranging from 10^0^ to 10^7^ gene copies. Analysis of qPCR data, including quantification and melt curve analysis, was done through the Bio-Rad CFX Manager Software (version 1.5). Efficiencies across all qPCR experiments ranged from 82.6-99.9%, with *R*^2^ values >0.99 for all standard curves. Final qPCR products were also verified using 1% agarose gels.

### Water chemistry

Water that was filtered through a 0.45-μm syringe filter prior to storage at -20°C was used for subsequent assays. Total ammonia, nitrite, and nitrate concentrations were measured in all samples using colourimetric assays following previously established protocols (9). The pH of water samples was measured using a LAQUAtwin pH-33 meter (Horiba Advanced Techno Co., Ltd.). Additionally, general water hardness (GH) and carbonate hardness (KH) were measured using the GH and KH Test Kit (Freshwater; API); alkalinity was measured using the MultiTest Marine pH & Alkalinity kit (Seachem).

### 16S rRNA gene sequencing and analysis

Primers 515F-Y and 926R were used to target the V4-V5 region of 16S rRNA genes (28, 29). For each sample, 25-μL PCR amplifications contained 1X ThermoPol buffer, 15 μg of bovine serum albumin, 200 μM of dNTPs, 0.2 μM of both forward and reverse primers, 0.625 units of Hot Start *Taq* DNA Polymerase (New England Biolabs, MA, USA), and 1-10 ng of template DNA. The PCR cycling conditions consisted of an initial denaturation at 95°C for 3 min, followed by 35 cycles of denaturation at 95°C for 30 s, annealing at 50°C for 30 s, and extension at 68°C for 1 min, with a final extension time of 7 min at 68°C. Individual samples were amplified in triplicate, with uniquely barcoded adapters attached to primers to allow sequencing of pooled samples. Additional no template and positive controls (1:1 *Aliivibrio fischeri* and *Thermus thermophilus* DNA) were included during PCR amplification. The resultant triplicate PCR products for each sample were pooled and quantified using a 1% agarose gel stained with GelRed (Biotium, CA, USA). Quantified pooled triplicates were then combined in equimolar concentrations based on gel quantification to create an amplicon library, which was sequenced (2 × 250 bases) using a MiSeq instrument (Illumina) following manufacturer’s protocols.

Following demultiplexing of paired-end sequence reads using MiSeq Reporter software version 2.5.0.5 (Illumina), analysis was done using QIIME2 (version 2020.6) implemented through the AXIOME3 pipeline (30). Quality trimming, primer sequence removal, denoising, paired-end sequence merging, chimera removal, and final generation of an amplicon sequence variant (ASV) table was done using DADA2 (31). Taxonomic classification of ASVs was done through AXIOME3 using the SILVA database release 138 (32). A phylogenetic tree of ASV sequences was generated using FastTree (33). Additional ASV table analysis was performed using the AXIOME3 pipeline, including rarefaction to the lowest sample count (8417) prior to calculation of beta diversity metrics (weighted UniFrac, Bray-Curtis) and generation of triplot ordinations. PERMANOVA testing was performed through QIIME2 (version 2020.6) using ‘beta-group-significance’ available through the ‘diversity’ plugin. All DNA sequences were deposited in the NCBI Sequence Read Archive under the BioProject accession number PRJNA780914.

## Results

### Aquarium samples

Aquarium biofilter and water samples (Table S1) were collected from 38 freshwater aquaria (FW-F01 – F38) and 8 saltwater aquaria (SW-F01 – F08). Samples were collected from both community members and retail pet stores from the Region of Waterloo and the city of Mississauga in Southwestern Ontario, Canada. Approximately half of the sampled aquaria contained live plants. The average number of fish across all aquaria was ∼22, ranging from 0-150. Several aquaria were populated with cichlids, algae eaters (e.g., “Plecos”), or guppies, and many contained mixed tropical or marine fish; one aquarium housed a turtle. Aquarium ages averaged 3 years, ranging from 1 month to 13 years, and were subject to diverse maintenance regiments (Table S1). Water used for most aquaria was sourced from municipal, reverse osmosis, distilled, or bottled water. Water changes occurred at frequencies ranging from once a week to once every eight weeks. For most aquarium biofilters, the sponge or floss material was never replaced; however, several owners replaced sponge/floss material as often as once a week to a frequency of every two years of use. At the time of aquarium establishment, five of the freshwater biofilters and one saltwater aquarium biofilter were inoculated with aquarium supplements. Within six months prior to sampling, only one freshwater aquarium and two saltwater aquaria had been exposed to antibiotics. Sampled aquaria covered a wide range of temperature (19.3-28.8°C), alkalinity (0-14), pH (6.2-9.3), general hardness (1-52 dGH), and carbonate hardness (1-38 dKH) (Table S2).

Measured concentrations of total ammonia in sampled aquaria were relatively low, with an average concentration of ∼59 μg/L NH_3_-N. Nitrite was usually undetected, or low in most aquaria, with only three aquaria with elevated levels of >1 mg/L NO_2_^-^-N (Table S2). In contrast, nitrate concentrations in most aquaria were relatively high (>1 mg/L NO_3_^-^-N).

### Aquarium biofilter microbial communities

A total of 12,241 unique amplicon sequence variants (ASVs) were identified in the 16S rRNA gene sequence data for 38 fresh and 8 saltwater samples, with a total of 969,251 reads and an average of 21,071 reads per sample. Both *Proteobacteria* and *Bacteroidota* dominated phylum-level ASV affiliation for all fresh and saltwater samples (Fig. 1). Members of the *Planctomycetota* were also consistently detected across all samples, ranging between 1-13% of all ASVs for each sample. Several additional phyla were present in most samples at >1%, including *Acidobacteria, Chloroflexi, Cyanobacteria, Myxococcota, Nitrospirota*, and *Verrucomicrobiota* (Fig. 1). Although sampled biofilter communities appeared similar at the phylum level, profiles were more distinct when displayed at the family level (>2%; Fig. 2). Members of the families *Saprospiraceae* and *Rhodobacteraceae* were present in both fresh and saltwater biofilters; *Rhodobacteraceae* were more abundant in saltwater samples (between 7-24%) than in freshwater samples (between 2-9%). Taxa from families *Cyclobacteraceae* and *Woeseiaceae* appeared only in saltwater biofilter samples at >2% abundance thresholds, whereas members of *Chitinophagaceae, Microscillaceae, Comamonadaceae*, and *Sphingomonadaceae* were consistently present for the majority of freshwater samples only above this same threshold (Fig. 2).

**Figure 1.**
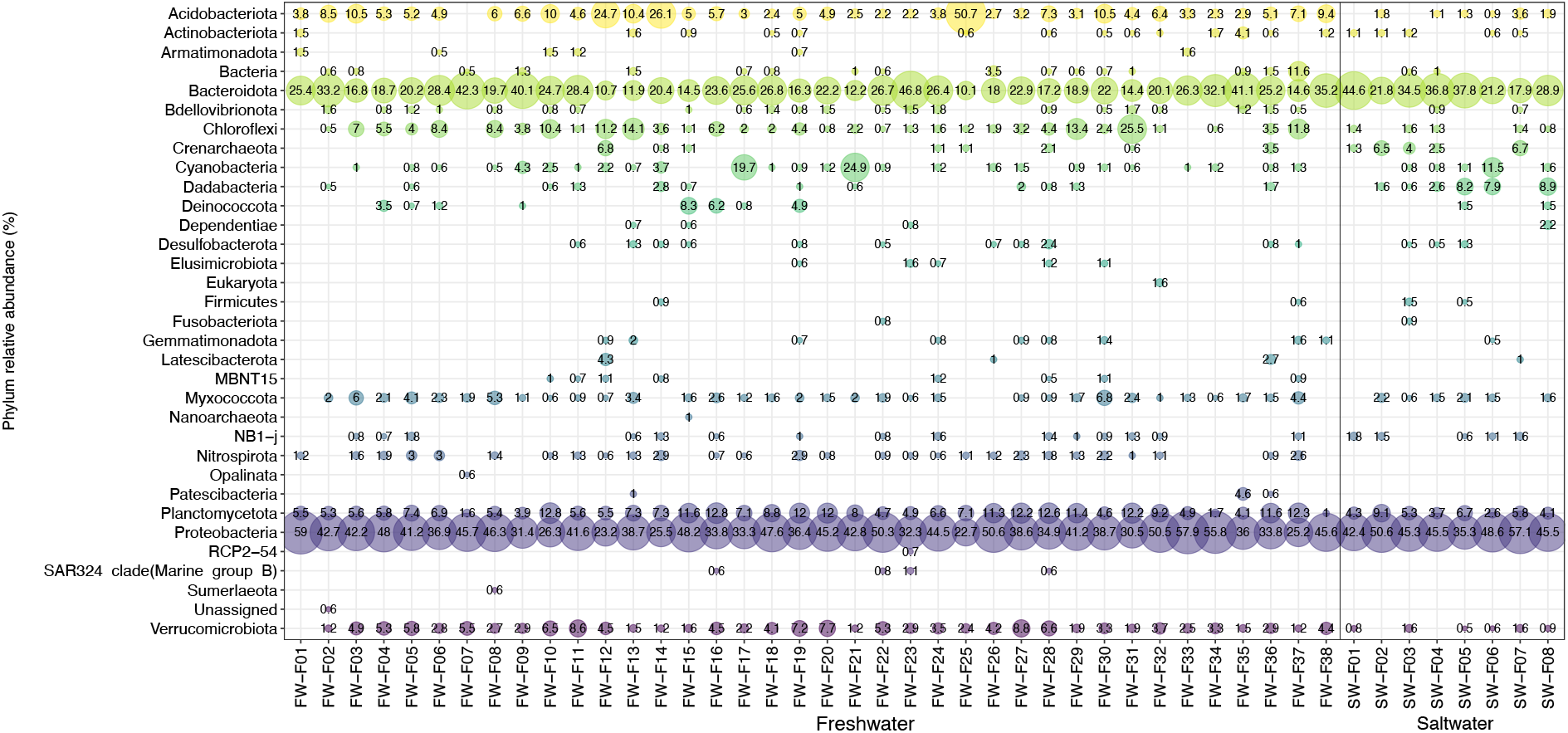
Relative abundance plot of 16S rRNA gene profiles of aquarium biofilter samples at the phylum level (>0.5%). Taxonomy has been assigned using SILVA 138 SSU database. Size of the bubble and the number represent the relative abundance of each phylum within each aquarium biofilter sample. Any phylum below 0.5% RA within a sample do not show on the plot.

**Figure 2.**
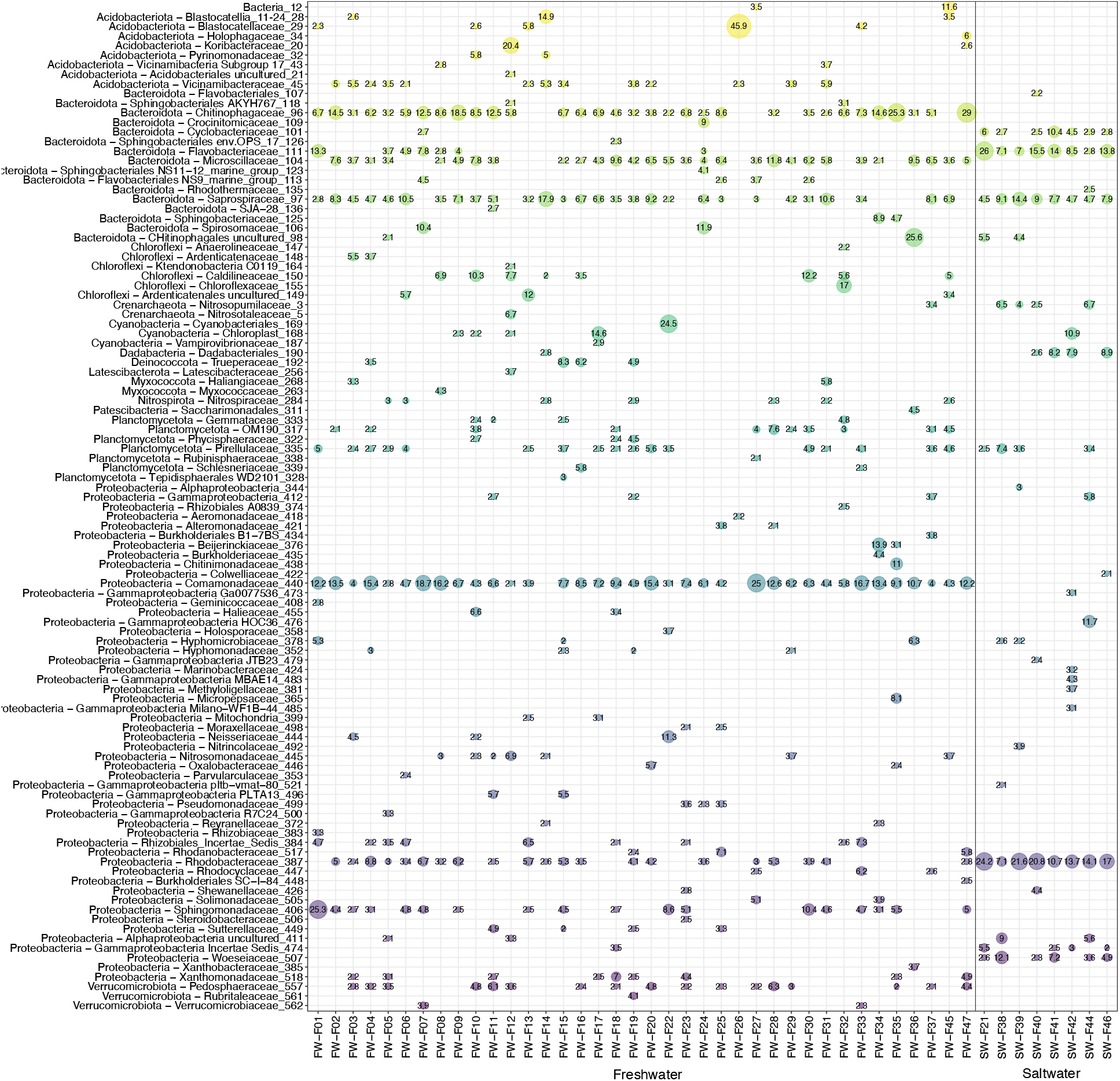
Relative abundance plot of 16S rRNA gene profiles of aquarium biofilter samples at the family level (> 2% RA). Taxonomy has been assigned using SILVA 138 SSU database, with taxonomy shown either at the family level of at the lowest assigned taxonomic level. Any samples containing taxa from a specific family that are below 2% RA are not shown.

Microbial community profiles of saltwater and freshwater samples were unique, grouping distinctly from one another in multidimensional space (Fig. 3A, pseudo-*F* = 9.57, *p* = 0.001). Although salinity explained the overall composition of sampled aquarium biofilters, other measured parameters also affected microbial community composition. Within freshwater aquarium biofilter samples specifically, correlation analysis revealed that aquarium age, general water hardness, and aquarium temperature all correlated significantly with 16S rRNA gene profiles (*R*^2^ > 0.3, *p* < 0.05, Fig. 3B). Several species (> 1% weighted average across samples) grouped in the centre of ordination space, indicating that these taxa are more commonly present across freshwater samples, including taxa affiliated with *Comamonadaceae, Vicinamibacteraceae, Rhodobacter*, and *Terimonas*. In contrast, *Blastocatellaceae*-associated taxa grouped away from the centre towards samples that had lower general hardness, indicating that this species was at a higher abundance in those samples compared with other samples in ordination space (Fig. 3B). When considering saltwater samples specifically, only aquarium size correlated with overall microbial community profiles (*R*^2^ > 0.3, *p* < 0.05), and this was also true for combined analysis with salt and freshwater samples. However, the majority of saltwater biofilter samples were collected from the same retail location, where the aquaria were relatively large compared to most at-home aquaria (Table S2). Only one other saltwater sample with a size much smaller in comparison could be included in the analysis because it had all associated metadata, whereas most freshwater aquaria were smaller, likely explaining why aquarium size was significantly correlated with community composition.

**Figure 3.**
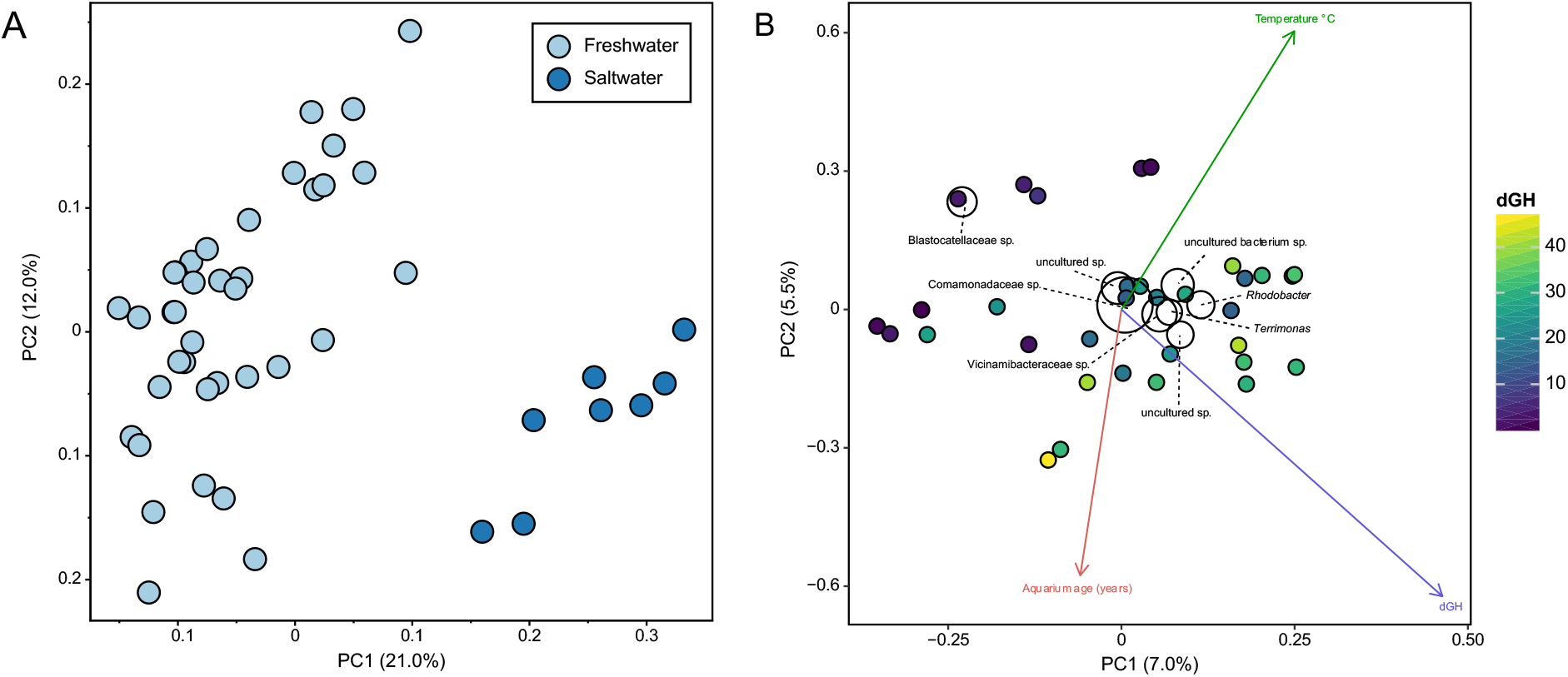
Principal coordinates and triplot analysis of 16S rRNA gene profiles of aquarium biofilters. Grouping between the saltwater and freshwater samples is shown in the PCoA plot (A) which was calculated using the weighted UniFrac metric on samples rarefied to the lowest sample count. An ordination based on the Bray-Curtis metric using only the freshwater samples (B) shows that the freshwater biofilm communities are correlated with temperature, aquarium age, and dGH (R^2^ > 0.3, *p* < 0.05). Taxa collapsed at the species level present at an average abundance of greater than 1% are displayed on the plot by the labelled black circles. Size of the circles illustrates the average relative abundance of the species across samples, while their placement reflects the correlation specific species have with different samples. The axes on both plots display the percent variation within the samples illustrated by each of the two principal components displayed.

### Ammonia oxidizer abundance in aquarium biofilters

Abundances of ammonia oxidizers were explored using qPCR to target the *amoA* gene of AOA, AOB, and comammox *Nitrospira*. Additionally, the 16S rRNA gene copies from bacteria and archaea were quantified. An initial screening with both primers targeting the *amoA* gene from the two different clades of *Nitrospira*, clade A and clade B (12), revealed that there were no detectable clade B comammox *Nitrospira* present in the aquarium biofilter samples (data not shown). Therefore, only primers associated with clade A comammox *Nitrospira* (comaA-244F(a-f) and comaA-659R(a-f)) were used for qPCR analysis.

Comammox *Nitrospira amoA* genes were detected across all 38 freshwater aquarium biofilter samples and were dominant ammonia oxidizers within 30 of the freshwater biofilters (Fig. 4). The *amoA* genes from AOA dominated seven of the freshwater biofilter samples. In contrast, AOB *amoA* genes dominated only one freshwater biofilter sample, and occurred at >1% (compared to comammox *Nitrospira* and AOA) in only five other freshwater biofilter samples (Fig. 4). Unlike freshwater aquarium biofilters, no comammox *Nitrospira amoA* genes were detected in any of the saltwater samples. For the eight sampled saltwater biofilters, AOA were the dominant ammonia oxidizers in five samples, AOB were dominant in two samples, and one saltwater sample had even proportions of AOA and AOB (Fig. 4).

**Figure 4.**
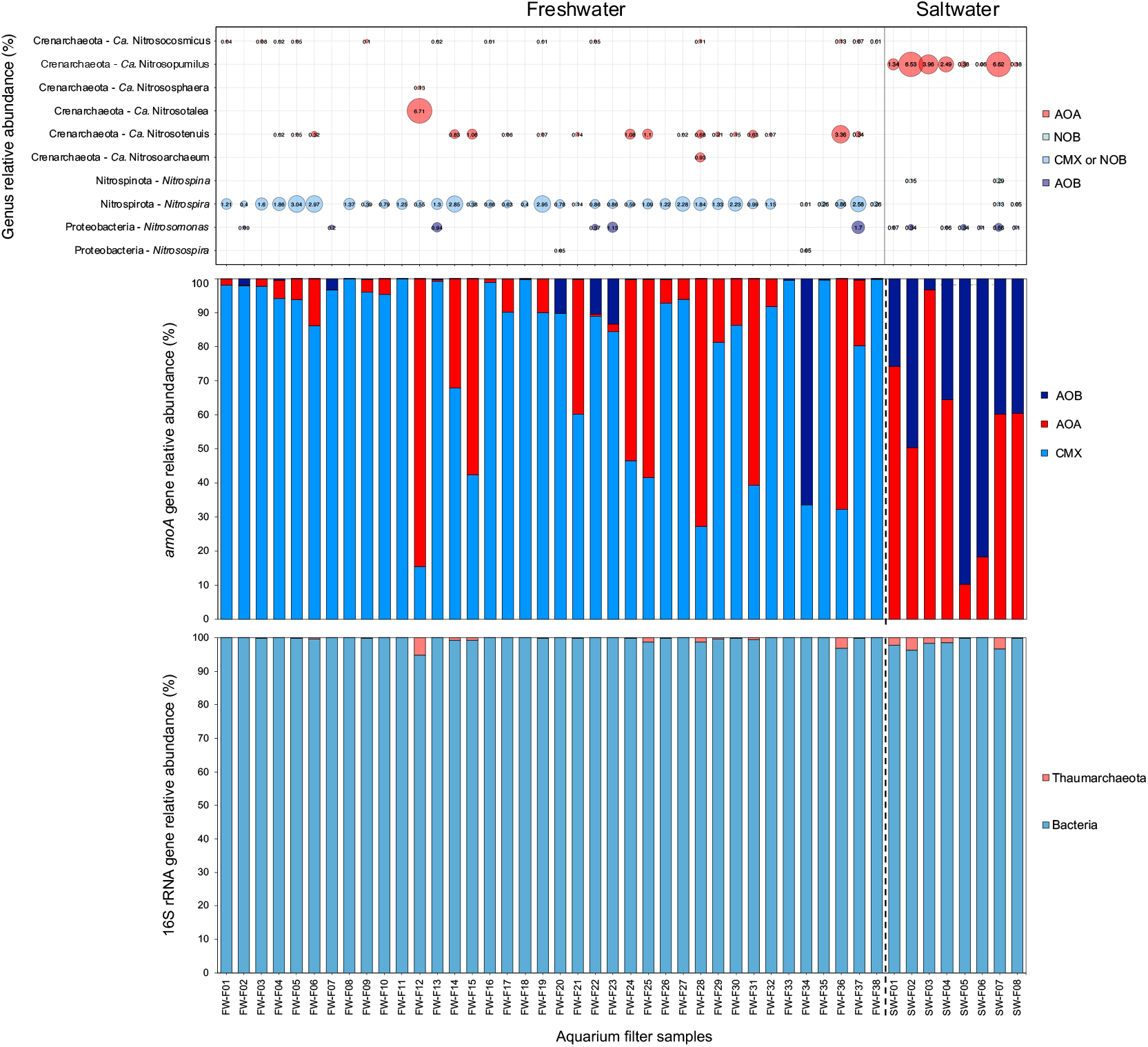
Abundance of microorganisms associated with ammonia oxidation were explored using both 16S rRNA gene profiles and gene abundances of the *amoA* gene, a marker for ammonia oxidation. The qPCR results of relative abundance of *amoA* genes from all three groups of ammonia oxidizers (AOA, AOB, and CMX) detected in biofilter samples are shown with the gene relative abundance of each group determined using the gene copy proportion for a single group relative to the total *amoA* gene copy number per ng of DNA from all three groups (middle). Relative gene abundance of 16S rRNA genes from bacteria and the archaeal phylum Thaumarchaeota or Thermoproteota, associated with AOA is shown on bottom. The genera associated with ammonia or nitrite oxidizing microorganisms present in the aquarium biofilter samples identified from the 16S rRNA gene sequences are illustrated by the bubble plot (top) with the size and number for each bubble representing the RA of each genus in each sample. Colours of bars and bubbles represent the respective groups of ammonia oxidizers, nitrite oxidizers, or comammox.

In addition to quantification of *amoA* gene copies, 16S rRNA genes associated with bacteria and archaea were quantified. All aquarium biofilter samples were dominated by bacterial 16S rRNA genes (>94%), whereas 16S rRNA genes associated with archaea were present at lower abundances across all samples, with some samples containing less than 2 copies/ng DNA (Table S5, Fig 4). Most samples that had higher relative abundances of 16S rRNA genes associated with archaea were also samples that had high relative abundances of AOA *amoA* genes (Fig. 4).

Using the 16S rRNA gene sequence data, we compared known nitrifiers to ammonia oxidizers quantified using qPCR (Fig. 4). For all saltwater samples, *Nitrosopumilus* was the only detected AOA. Detected genera of known AOB included *Nitrosomonas* and these were found in all but one sample. Based only on 16S rRNA gene sequencing, the NOB in the saltwater samples included the genus *Nitrospina* and *Nitrospira*, which were likely strict nitrite oxidizers because no comammox *Nitrospira* were detected in the saltwater samples. However, not all saltwater samples contained sequences affiliated with known NOB genera (e.g., SW-1, SW-3 – SW-5). In contrast to the saltwater samples, freshwater samples had a higher diversity of AOA, including *Nitrosocosmicus* and *Nitrosotenuis*, which occurred in many samples. The AOA genera *Nitrososphaera, Nitrosotalea*, and *Nitrosoarchaeum* were found in only one sample each. Two genera of AOB, *Nitrosomonas* and *Nitrosospira*, were detected in several freshwater samples. The genus *Nitrospira* was detected across all but two freshwater samples and represent both comammox *Nitrospira* and canonical NOB *Nitrospira*, which are indistinguishable using 16S rRNA genes. No other NOB-associated genera were detected in the freshwater samples.

Overall, relative abundances of ammonia oxidizers detected with 16S rRNA gene sequencing matched qPCR relative abundances well, although there were a few discrepancies. For example, samples FW-F07 and FW-F33 did not have any detectable *Nitrospira* via 16S rRNA gene analysis but did show detectable comammox *Nitrospira* with *amoA* gene qPCR. Additionally, FW-F33 did not have any genera associated with known nitrifiers from 16S rRNA gene data. Because there were several ASVs with taxonomic assignments at the family level to families associated with nitrification (e.g., *Nitrosomonadaceae*), these ASVs may be associated with poorly characterized nitrifiers.

Along with determining the dominant ammonia oxidizers present in the biofilters, we explored correlations between ammonia oxidizer abundance and aquarium characteristics. Our results indicated that total ammonia concentration was likely linked to ammonia oxidizer relative abundances. For the samples collected, there was only one aquarium that had a relatively high concentration of total ammonia (FW-F34, 650.6 μg/L NH_3_-N) and this sample also had the highest relative abundance of AOB based on *amoA* gene quantification (Fig. 4).

## Discussion

### Aquarium biofilter microbial communities

We show that aquarium biofilters possess distinct microbial communities in salt and freshwater environments at both the level of total community and also at the level of nitrifier-associated genera, reinforcing salinity as an important driver of microbial community structure in aquarium biofilters. The 16S rRNA gene sequencing data show that the phyla *Proteobacteria, Bacteroidota*, and *Planctomycetota* dominated all biofilters, which was also observed in a previous study of freshwater aquarium water (34). The presence of these phyla, along with the others observed in our microbial community data, were also observed for the microbial communities of RAS biofilters (35, 36). At the phylum level, there was little difference between the salt and freshwater communities and the relative abundances were consistent among samples, indicating a common microbial community composition among aquaria. Although differences in microbial communities between the two environments was evident at lower taxonomic levels (e.g., family level).

Differentiation in microbial community composition at a functional level between salt and freshwater was also notable within the nitrifiers, not only with the presence and absence of comammox *Nitrospira* in fresh and saltwater aquaria respectively, but in the AOA, AOB, and NOB present in the biofilter communities. The saltwater AOA were dominated by members from the genus *Nitrosopumilus*, which were originally discovered in marine aquarium gravel (18), while many freshwater aquaria contain AOA from the genera *Nitrosotenuis* and *Nitrosocosmicus*. Differences in nitrifiers between salt and freshwater were also observed in past aquarium surveys (21). Within the freshwater AOA ASVs identified, some belonged to both *Ca*. Nitrosotenuis aquarius and *Ca*. Nitrosocosmicus hydrocola, which were discovered in a freshwater aquarium biofilter and a tertiary wastewater treatment system respectively (37, 38). There were only a few saltwater samples with detectable NOB genera either belonging to *Nitrospira* or *Nitrospina* spp. and they were present at quite a low relative abundance (< 0.5%) within the community. However, there was detectable nitrite within almost all the saltwater tanks sampled (Table S2), so perhaps the nitrite oxidizers were not well established within the biofilter material sampled.

Microbial profiles of freshwater biofilters revealed that community composition was correlated with temperature, aquarium size, and general hardness of the water. Other studies have demonstrated the importance of environmental, biological, and physical factors, such as temperature, filter support material, and fish species as being involved in differentiating biofilter microbial communities in systems like RAS, water treatment, and aquaponics (35, 39, 40). Although there is interest in the nitrifying members of biofilter communities for their role in converting toxic nitrogenous waste in aquarium systems, other microbial members also play important roles as heterotrophic organisms in the breakdown of organic waste products (e.g. dissolved organic matter), with the possibility to have members of the microbial community also capable of processes like denitrification, anaerobic ammonium oxidation, methanogenesis, and dissimilatory nitrate reduction to ammonia (DNRA) (36). The occurrence of these microbiological processes is dependent on the conditions within the biofilter and the microbial community composition, which highlights the importance of increasing our understanding of the ecology of aquarium biofilters to optimize the functionality of the microbial communities of aquarium biofilters, both in home aquaria and commercial systems.

### Abundance of ammonia oxidizers within aquarium biofilters

Our reassessment of the ammonia oxidizing microorganisms in aquarium biofilters revealed an unexpected abundance of comammox *Nitrospira* in freshwater aquarium biofilters, which revises a previous description of aquarium nitrification that focused on AOA (21). Both qPCR and 16S rRNA gene sequencing data showed that comammox *Nitrospira* and AOA together dominate as ammonia oxidizers in freshwater aquarium biofilters, whereas saltwater aquarium biofilters are dominated by either AOA or AOB. Results from qPCR and 16S rRNA gene sequencing were consistent for most samples showing similar abundances of AOA based on both *amoA* and 16S rRNA gene copy numbers, and relative abundances of AOA-associated genera. Although AOA *amoA* gene abundances were similar to those of archaeal 16S rRNA genes, 16S rRNA gene abundances exceeded corresponding functional gene abundances for some samples. This was also observed in a previous survey of aquarium oxidizers (21), and it may be that not all taxa targeted by the *Thermoproteota* primer set correspond to AOA. The presence of *Nitrospira* via 16S rRNA gene sequencing was detected in all but two freshwater samples. Considering that comammox *Nitrospira* were detected via *amoA* gene qPCR, the absence of *Nitrospira* in the 16S rRNA gene sequencing results for those samples was unexpected. There were no other ASVs assigned to the phylum *Nitrospirota* present in these two samples that may have represent the comammox *Nitrospira* detected via qPCR. Additionally, the level of *amoA* gene copies associated with comammox *Nitrospira* were not unusually low compared with other samples, making their absence in the 16S rRNA gene sequencing data difficult to explain by low abundance within the sample. Some insight from future *amoA* sequencing data may help to clarify the lack of *Nitrospira* ASVs detected via 16S rRNA gene sequencing in these two samples.

Overall, the ubiquitous presence of comammox *Nitrospira* within freshwater aquarium biofilters is consistent with their presence in similar environments, such as recirculating aquaculture system (RAS) biofilters, groundwater-fed biofilters, and aquaponics systems (13, 23, 41). Relatively low ammonia concentrations and high surface area for biofilm growth within biofilters are ideal conditions for microorganisms with slow growth rates and high yields, as originally predicted for comammox *Nitrospira* (6). Additionally, the absence of comammox *Nitrospira* in saltwater aquarium biofilters reflects results from other studies showing that comammox *Nitrospira* are not found within marine environments. Although comammox *Nitrospira* have been identified in brackish, salt marsh, and estuarine environments (42, 43), it seems that the role of autotrophic ammonia oxidation is occupied by AOA and AOB in fully marine environments, including home marine aquaria.

Only comammox *Nitrospira* from clade A were associated with freshwater biofilters. In previous studies, most comammox *Nitrospira* associated with wastewater treatment systems (e.g., activated sludge and rotating biological contactors) also belong to clade A *Nitrospira* (9, 44–47). In contrast, clade B comammox *Nitrospira* have been identified frequently as the dominant comammox bacteria in forest soils, paddy soils, plateau soils, and river sediments, and have been experimentally confirmed as active contributors to soil nitrification in forest soils (45, 48). Often the presence of clade B members is accompanied by clade A comammox *Nitrospira*, as was identified in metagenomic surveys of terrestrial subsurface samples (49). Although most studies examining water treatment systems have identified clade A comammox, clade B dominated a groundwater-fed rapid sand filter existing alongside other clade A comammox *Nitrospira* (13). Factors influencing distributions of these two clades are unclear; however, based on our results, clade A comammox *Nitrospira* dominate freshwater aquarium biofilters.

### Niche differentiation of ammonia oxidizers in aquarium biofilters

We observed that the majority of the aquarium biofilters were dominated by either AOA or comammox *Nitrospira*, although it is not clear what factors may be contributing to their distributions. Previously, pH and ammonia concentration were identified as two key factors governing the abundance of different ammonia oxidizing microorganisms in the environment (50). Our results do support the observations from previous studies that comammox *Nitrospira* and AOA are dominant in lower ammonia conditions. Previous work shows that both AOA and comammox *Nitrospira* compete favourably in low ammonia conditions, with several cultivated representatives having high affinities for ammonia including *Nitrospira inopinata, Ca*. N. kreftii, *Nitrosopumilus maritimus*, and *Ca*. Nitrosotenuis aquarius (38, 51, 52). However, at this time there are only have a few cultivated representatives of comammox *Nitrospira* where ammonia affinities have been measured. Additionally, there are several AOA species that are also known to have tolerance for higher ammonia concentrations, such as those from the genus *Nitrosocosmicus* (53). We noticed that there was a higher relative abundance of *Nitrosotenuis* spp. than *Nitrosocosmicus* spp. for freshwater samples.

Although most freshwater aquaria sampled were low in ammonia, there was one sampled aquarium (FW-F34) that had a high concentration of total ammonia and was dominated by AOB *amoA* genes. In all other freshwater samples, AOB fell below 14% RA, with most being < 1% of the total ammonia oxidizers detected with qPCR. The low abundance of both detected comammox *Nitrospira* and AOB is reflected in the overall microbial community of FW-F34 where associated genera of *Nitrospira* and *Nitrosomonas* are both present at a RA of > 0.5%. Although this sample suggests AOB might favour high ammonia concentrations over AOA and comammox, more data is needed to support this hypothesis. The effects of ammonia concentration on ammonia oxidizer abundance may be easier to test under experimental conditions with controlled ammonia concentrations as most aquaria naturally have low levels of ammonia with well-established nitrifying communities in their biofilms.

The presence of AOA and comammox *Nitrospira* identified via genes for ammonia oxidation suggests that they are performing ammonia oxidation in these systems; however, both have the potential for alternative metabolisms. Some AOAs have the potential for mixotrophy, while some comammox *Nitrospira* have the genetic capabilities to utilize different forms of nitrogen such as urea and cyanate (9, 38, 54). With the potential for alternative metabolisms, it is important to confirm the ammonia oxidation activity of AOA and comammox with addition activity-based experiments. Additional factors that were not explored in this study that may contribute to niche differentiation between AOA and comammox include dissolved oxygen (DO) concentrations, which have been noted in other studies where comammox *Nitrospira* are found at high abundance in environments with low DO concentrations (44, 47).

Overall, this work has further clarified our understanding of ammonia oxidation in aquarium biofilters revealing that comammox *Nitrospira* are ubiquitous in freshwater biofilters, and the dominant ammonia oxidizers are either comammox *Nitrospira* and AOA. Now that we hopefully have completed our picture of the autotrophic microbial members involved in aquarium ammonia oxidation, future research should work to further address the factors that may be involved in niche differentiation in these environments and explore the contributions from each group to ammonia oxidation in the biofilters. An improved understanding of this microbially mediated process in aquaria is important to improve and optimize current water treatment in the aquaculture industry and could potentially lead to the development of new and improved supplements for aquarium biofilters.

## Acknowledgements

We would like to thank members of the Kitchener-Waterloo Aquarium Society and Big Al’s Aquarium Supercentre in Kitchener, ON for their participation in this study and providing us with aquarium filter and water samples. We also acknowledge that most aquarium samples were collected from within the Haldimand Tract, which is land that was granted to the Haudenosaunee of the Six Nations of the Grand River, and within the territory of the Neutral, Anishinaabe, and Haudenosaunee peoples. We thank Rachel Beaver, Katja Engel, and Alex Umbach for assistance with sequencing. This work was supported by a Postgraduate Scholarship to MMM and a Discovery Grant to JDN, both from the National Science and Engineering Research Council of Canada (NSERC).

## Notes

### Competing Interest Statement

The authors have declared no competing interest.

