## Supplementary Information for "Microbial community analysis of biofilters reveals a dominance of either comammox *Nitrospira* or archaea as ammonia oxidizers in freshwater aquaria"

Michelle M. McKnight^1^ and Josh D. Neufeld^1^

^1^Department of Biology, University of Waterloo, Waterloo, Ontario, Canada

**qPCR methods**

*Generation of qPCR gene standards*

DNA templates used to generate standard curves for the comammox *amoA*, AOA *amoA*, AOB *amoA*, and 16S rRNA gene thaumarcheotal qPCR were generated using PCR with the same primers described in the main manuscript. Following PCR amplification of these target sequences, PCR products were purified using the Wizard SV Gel and PCR Clean-Up System (Promega). For the 16S rRNA gene qPCR standard, the target gene was amplified from a pUC57-Kan vector containing the target 16S rRNA gene fragment from *Thermus thermophilus* flanked by M13 primers. The M13 primers were used to amplify the 16S rRNA gene target and following PCR amplification the PCR product was also purified as described for the other four DNA template targets. Standards were stored in aliquots at concentrations of around 10^10^ copies/μL at -20°C. The DNA concentration of standard aliquots were measured on the same day the qPCR was performed using a Qubit dsDNA HS Assay Kit (ThermoFisher Scientific). Measurements were done in duplicate using 4 μL of sample, and the average concentration was used to determine the copy number concentrations for the qPCR standard curves alongside the molecular mass of each standard amplicon sequence as described below.

For the AOA, AOB, bacterial, and archaeal targets, exact sequences of the amplicons were known, and these sequences were used to determine the exact copy numbers added to each qPCR reaction based on the molecular weight of the fragment (Table S1-S2). Since the standard for clade A comammox *Nitrospira* was amplified from a mixture of different comammox *amoA* sequences from aquaria, a molecular weight of an exact sequence could not be used to determine the copy number. Therefore, the average molar mass per bp of 650 g/mol/bp was used to determine the copy number concentrations added to the qPCR reactions. Specifically, the ThermoFisher Sci DNA copy number and dilution calculator (<https://www.thermofisher.com/ca/en/home/brands/thermo-scientific/molecular-biology/molecular-biology-learning-center/molecular-biology-resource-library/thermo-scientific-web-tools/dna-copy-number-calculator.html>) was used to determine copy number based on the concentration of the standard DNA template solution. For the other standards with known DNA sequences, the Science Primer copy number calculator was used to determine copy number based off exact molecular weight of the fragment and DNA concentration (<http://www.scienceprimer.com/copy-number-calculator-for-realtime-pcr>).

*Preparation of qPCR standard curve dilution series*

The standard curve DNA samples were prepared using a freshly thawed standard aliquot that had a concentration of ~10^10^ copies/μL. Standard dilutions were prepared in a 10-fold dilution series from magnitudes of 10^9^ copies/μL to 10^0^ copies/μL in 100 μL volumes. Each dilution was made with using 10 mM Tris-HCl buffer containing 0.05% Tween-20 to minimize adherence of any DNA to the tube.

**Table S5.** Gene copies for both *amoA* and 16S rRNA gene targeted qPCRs for each sample. Copies are expressed per ng of extracted DNA.

| Sample ID | Sampling date | Location | City | Aquarium type | Filter material | Live plants (y/n) | Added supplements (y/n) | Antibiotic treatment  (< 6 mo.) | Water source | Fish type | Aquarium age (years) | Fish | Last sponge replacement (months) | Frequency of water changes (weeks) |
| --- | --- | --- | --- | --- | --- | --- | --- | --- | --- | --- | --- | --- | --- | --- |
| FW-F01 | 03/05/2019 | Residential 1 | Waterloo | Fresh | Sponge | Yes | No | No | Tap water | Discus, angel | 7 | 12 | 12 | 1 |
| FW-F02 | 03/05/2019 | Residential 2 | Kitchener | Fresh | Sponge | Yes | No | No | Tap water | Guppies | 0.5 | 10 | Never | 1 |
| FW-F03 | 03/05/2019 | Residential 3 | Guelph | Fresh | Sponge | No | No | No | Tap water | Cichlids, pleco | 3 | 37 | 12 | 4 |
| FW-F04 | 03/05/2019 | Residential 3 | Guelph | Fresh | Sponge | Yes | No | No | Tap water | Mixed tropical | 2 | 67 | 12 | 4 |
| FW-F05 | 03/05/2019 | Residential 4 | Drayton | Fresh | Sponge | No | No | No | Tap water | Cichlids, petricola | 9 | 10 | 60 | 1 |
| FW-F06 | 03/05/2019 | Residential 4 | Drayton | Fresh | Sponge | No | No | No | Tap water | Cichlid | 9 | 5 | 60 | 1 |
| FW-F07 | 03/05/2019 | Residential 5 | Drayton | Fresh | Sponge | No | No | No | Tap water | Guppies, pleco | 0.1 | 8 | 1 | 4 |
| FW-F08 | 03/05/2019 | Residential 5 | Waterloo | Fresh | Sponge | Yes | No | No | Tap water | Guppies, pleco | 2 | 20 | Never | 2 |
| FW-F09 | 03/05/2019 | Residential 5 | Waterloo | Fresh | Sponge | Yes | No | No | Tap water | Barbs, pleco | 1 | 13 | Never | 4 |
| FW-F10 | 03/05/2019 | Residential 6 | Fergus | Fresh | Floss | Yes | No | No | Tap water | Cichlid, pleco | 0.5 | 2 | Never | 8 |
| FW-F11 | 03/05/2019 | Residential 6 | Fergus | Fresh | Floss | Yes | Yes | Yes | Tap water | Mixed tropical | 1 | 25 | Never | 4 |
| FW-F12 | 03/09/2019 | Residential 7 | Mississauga | Fresh | Floss | No | No | No | Tap water | Cichlid | 0.5 | 1 | 1 | 5 |
| FW-F13 | 03/09/2019 | Residential 7 | Mississauga | Fresh | Sponge | No | No | No | Tap water | Turtle | 9 | 1 | 1 | 4 |
| FW-F14 | 03/18/2019 | Residential 8 | Waterloo | Fresh | Sponge | No | No | No | Tap water | Cichlid | 12 | 3 | 8 | 4 |
| FW-F15 | 04/02/2019 | Residential 9 | Drayton | Fresh | Floss | Yes | No | No | Tap water | Mixed tropical | 3 | 20 | 6 | 4 |
| FW-F16 | 04/02/2019 | Residential 5 | Drayton | Fresh | Sponge | Yes | No | No | Tap water | Guppies, plecos | 0.3 | 30 | 4 | 4 |
| FW-F17 | 04/02/2019 | Residential 10 | Kitchener | Fresh | Floss | No | No | No | Tap water | Endlers, crayfish | 4 | 20 | 24 | 2 |
| FW-F18 | 04/02/2019 | Residential 11 | Cambridge | Fresh | Floss | Yes | Yes | No | Tap water | Plecos | 0.75 | 10 | 1 | 1 |
| FW-F19 | 04/02/2019 | Residential 11 | Cambridge | Fresh | Floss | No | Yes | No | Tap water | Cichlids | 0.3 | 10 | 1 | 1 |
| FW-F20 | 04/02/2019 | Residential 12 | Kitchener | Fresh | Sponge | Yes | Yes | No | Tap water | Cichlids | 6 | 12 | Never | 2 |
| FW-F21 | 04/02/2019 | Residential 14 | Guelph | Fresh | Sponge | Yes | No | No | Tap water | Danios | 0.75 | 4 | Never | 4 |
| FW-F22 | 04/02/2019 | Residential 15 | Waterloo | Fresh | Sponge | No | No | No | Tap water | Unknown | 2 | Unknown | Unknown | Unknown |
| FW-F23 | 04/02/2019 | Residential 15 | Waterloo | Fresh | Sponge | No | No | No | Tap water | Unknown | 2 | Unknown | Unknown | Unknown |
| FW-F24 | 04/02/2019 | Residential 15 | Waterloo | Fresh | Sponge | No | No | No | Tap water | Unknown | 2 | Unknown | Unknown | Unknown |
| FW-F25 | 04/02/2019 | Residential 16 | Guelph | Fresh | Sponge | Yes | No | No | Tap water | Mixed tropical | 0.5 | 75 | Never | 1 |
| FW-F26 | 04/02/2019 | Residential 16 | Guelph | Fresh | Sponge | Yes | No | No | Tap water | Cichlids | Unknown | 5 | 2 | 1 |
| FW-F27 | 04/02/2019 | Residential 16 | Guelph | Fresh | Sponge | No | No | No | Tap water | Cichlids | 0.5 | 30 | Never | 1 |
| FW-F28 | 04/02/2019 | Residential 16 | Guelph | Fresh | Sponge | No | No | No | Tap water | Mixed tropical | 1 | 25 | Never | 1 |
| FW-F29 | 04/02/2019 | Residential 3 | Guelph | Fresh | Sponge | Yes | No | No | Tap water | Cichlids | 1 | 103 | 9 | 4 |
| FW-F30 | 04/02/2019 | Residential 3 | Guelph | Fresh | Floss | No | No | No | Tap water | Cichlids, plecos | 1 | 40 | 5 | 4 |
| FW-F31 | 04/02/2019 | Residential 17 | Waterloo | Fresh | Sponge | Yes | Yes | No | Ground and RO water | Mixed tropical | 3 | 50 | Never | 4 |
| FW-F32 | 04/02/2019 | Residential 18 | Waterloo | Fresh | Sponge | No | No | No | Tap water | Guppies, tetras | 2 | 12 | Never | 2 |
| FW-F33 | 04/02/2019 | Residential 18 | Waterloo | Fresh | Sponge | Yes | No | No | Bottled water | *Microctenopoma* | 0.5 | 2 | Never | 3 |
| FW-F34 | 04/02/2019 | Residential 18 | Waterloo | Fresh | Sponge | No | No | No | Bottled water | Mixed tropical | 1 | 30 | Never | 3 |
| FW-F35 | 04/02/2019 | Residential 18 | Waterloo | Fresh | Sponge | Yes | No | No | Tap and distilled water | *Cynodorichthys* | 0.5 | 2 | Never | 3 |
| FW-F36 | 05/02/2019 | Residential 19 | Kitchener | Fresh | Sponge | Yes | No | No | RO water | Tetras, raboras | 13 | 8 | 6 | 3 |
| FW-F37 | 05/12/2019 | Residential 7 | Mississauga | Fresh | Sponge | No | No | No | Tap water | Turtle | 9 | 1 | 1 | 4 |
| FW-F38 | 06/05/2019 | Residential 22 | Unknown | Fresh | Sponge | Unknown | Unknown | Unknown | Unknown | Unknown | Unknown | Unknown | Unknown | Unknown |
| SW-F01 | 04/02/2019 | Residential 13 | Guelph | Salt | Floss | Yes | No | No | Tap water | Damsels, clownfish | 3 | 8 | 4 | 4 |
| SW-F02 | 05/13/2019 | Residential 20 | Kitchener | Salt | Sponge | Yes | No | No | Tap water | Mixed marine | 13 | 12 | Never | 8 |
| SW-F03 | 05/17/2019 | Retail 1 | Kitchener | Salt | Floss | Yes | No | No | RO water | Mixed marine | 1.5 | 10 | 0.25 | 1 |
| SW-F04 | 05/17/2019 | Retail 1 | Kitchener | Salt | Floss | Yes | No | No | Tap water | Mixed marine | 1.5 | 12 | 0.25 | 1 |
| SW-F05 | 05/17/2019 | Retail 1 | Kitchener | Salt | Floss | No | No | Yes | Tap water | Tangs, wrasse | 1.5 | 3 | 0.25 | 1 |
| SW-F06 | 05/17/2019 | Retail 1 | Kitchener | Salt | Floss | No | Yes | Yes | Tap water | Mixed marine | 0.5 | 150 | 0.25 | 1 |
| SW-F07 | 05/17/2019 | Retail 1 | Kitchener | Salt | Floss | Unknown | Unknown | Unknown | Unknown | None | Unknown | Unknown | Never | 1 |
| SW-F08 | 05/21/2019 | Residential 21 | Unknown | Salt | Floss | Unknown | Unknown | Unknown | Unknown | Unknown | Unknown | Unknown | Unknown | Unknown |

**Table S1.** Summary of aquarium sample metadata, including information on aquarium type, maintenance, location, number of fish and sampling dates.

**Table S2.** Water chemistry, temperature, and aquarium size data for all aquarium biofilter samples collected.

| ID | Size (gallons) | Temperature (ºC) | pH | Alkalinity (meq/L) | dGH | dKH | Total NH_3_-N (μg/L) | NO_2_^-^-N (μg/L) | NO_3_^-^-N (μg/L) |
| --- | --- | --- | --- | --- | --- | --- | --- | --- | --- |
| FW-F01 | 120 | 28.0 | 8.6 | 14.0 | 3 | 20 | 16.7 | 21.6 | 2394.0 |
| FW-F02 | 15 | 25.0 | 8.1 | 14.0 | 21 | 10 | 0.0 | 0.0 | 4274.5 |
| FW-F03 | 55 | 22.0 | 8.0 | 4.5 | 30 | 7 | 0.0 | 19.9 | 27829.2 |
| FW-F04 | 29 | 24.0 | 8.0 | 2.5 | 31 | 7 | 12.5 | 45.1 | 39762.7 |
| FW-F05 | 30 | 24.4 | 8.3 | 3.5 | 32 | 9 | 0.0 | 0.0 | 4127.9 |
| FW-F06 | 20 | 24.4 | 8.2 | 3.0 | 32 | 7 | 0.0 | 14.1 | 5097.5 |
| FW-F07 | 10 | 25.6 | 8.2 | 4.0 | 14 | 11 | 16.7 | 2013.6 | 2262.0 |
| FW-F08 | 20 | 24.4 | 8.1 | 4.5 | 16 | 13 | 3.7 | 13.2 | 6225.1 |
| FW-F09 | 30 | 24.4 | 8.2 | 4.0 | 15 | 11 | 20.3 | 0.0 | 7403.5 |
| FW-F10 | 29 | 25.6 | 8.5 | 7.5 | 3 | 18 | 0.0 | 0.0 | 30337.0 |
| FW-F11 | 75 | 25.0 | 8.4 | 5.0 | 2 | 13 | 0.3 | 0.0 | 17516.9 |
| FW-F12 | 90 | 23.9 | 6.8 | 0.0 | 27 | 1 | 1.8 | 16.3 | 21473.1 |
| FW-F13 | 5 | 20.0 | 8.4 | 5.0 | 31 | 15 | 232.5 | 94.7 | 4425.8 |
| FW-F14 | 110 | 26.7 | 8.3 | 3.0 | 24 | 11 | 0.0 | 0.0 | 4313.0 |
| FW-F15 | 50 | 23.9 | 8.7 | 7.5 | 25 | 20 | 25.8 | 0.0 | 9535.9 |
| FW-F16 | 50 | 25.6 | 8.6 | 4.5 | 18 | 12 | 0.0 | 20.3 | 8203.7 |
| FW-F17 | 5 | 20.0 | 8.1 | 1.5 | 25 | 10 | 12.0 | 0.0 | 5149.2 |
| FW-F18 | 65 | 26.0 | 8.1 | 2.5 | 31 | 10 | 0.0 | 18.0 | 3364.0 |
| FW-F19 | 90 | 26.0 | 8.3 | 4.0 | 33 | 12 | 0.7 | 15.7 | 8223.9 |
| FW-F20 | 110 | 25.6 | 8.1 | 2.5 | 28 | 9 | 22.9 | 0.0 | 2756.5 |
| FW-F21 | 5 | 25.0 | 9.3 | 3.5 | 17 | 7 | 0.0 | 0.0 | 15.3 |
| FW-F22 | unknown | 27.7 | 7.5 | 2.0 | 52 | 4 | 250.8 | 233.0 | 106676.6 |
| FW-F23 | unknown | 27.7 | 8.3 | 7.5 | 46 | 17 | 109.0 | 246.2 | 37921.3 |
| FW-F24 | unknown | 27.7 | 8.1 | 4.0 | 38 | 8 | 70.6 | 46.8 | 46290.2 |
| FW-F25 | 125 | 27.0 | 8.6 | 12.0 | 4 | 28 | 59.7 | 15.0 | 14986.4 |
| FW-F26 | 15 | 22.0 | 8.6 | 10.5 | 3 | 23 | 14.9 | 0.0 | 1004.8 |
| FW-F27 | 20 | 28.8 | 8.6 | 16.5 | 5 | 38 | 16.6 | 0.0 | 90721.8 |
| FW-F28 | 155 | 27.3 | 8.1 | 4.5 | 8 | 13 | 54.2 | 0.0 | 34680.7 |
| FW-F29 | 29 | 26.7 | 8.2 | 4.0 | 39 | 12 | 0.0 | 0.0 | 1976.2 |
| FW-F30 | 75 | 23.9 | 8.3 | 4.0 | 41 | 9 | 0.0 | 17.1 | 27163.0 |
| FW-F31 | 46 | 23.9 | 7.9 | 1.5 | 19 | 3 | 0.0 | 0.0 | 6683.2 |
| FW-F32 | 10 | 19.3 | 8.0 | 4.0 | 17 | 10 | 0.0 | 0.0 | 899.5 |
| FW-F33 | 10 | 19.3 | 8.2 | 0.5 | 2 | 1 | 0.0 | 0.0 | 62.9 |
| FW-F34 | 10 | 19.5 | 6.2 | 0.5 | 4 | 1 | 650.6 | 0.0 | 4599.3 |
| FW-F35 | 10 | 19.7 | 6.9 | 0.5 | 1 | 2 | 0.0 | 0.0 | 849.5 |
| FW-F36 | 10 | 25.0 | 7.1 | 0.5 | 40 | 3 | 0.0 | 0.0 | 1012.2 |
| FW-F37 | 5 | 20.0 | 8.6 | 7.0 | 46 | 18 | 237.8 | 40.1 | 4943.2 |
| FW-F38 | unknown | unknown | 6.5 | 1.5 | 36 | 4 | 136.9 | 1029.4 | 60340.8 |
| SW-F01 | 55 | 22.6 | 8.0 | 4.0 | n/a | 11 | 16.2 | 0.0 | 24861.7 |
| SW-F02 | 180 | 25.6 | 8.0 | 5.0 | n/a | 14 | 0.0 | 42.4 | 59595.8 |
| SW-F03 | 240 | 25.6 | 8.1 | 4.5 | n/a | 12 | 0.0 | 42.4 | 2365.4 |
| SW-F04 | 280 | 25.6 | 8.1 | 8.0 | n/a | 22 | 299.8 | 221.5 | 9156.8 |
| SW-F05 | 170 | 25.6 | 8.3 | 6.5 | n/a | 16 | 8.2 | 0.0 | 21177.8 |
| SW-F06 | 200 | 25.6 | 8.2 | 4.5 | n/a | 13 | 45.4 | 14.0 | 4082.3 |
| SW-F07 | unknown | 25.6 | 8.1 | 5.5 | n/a | 15 | 12.7 | 45.2 | 3412.8 |
| SW-F08 | unknown | unknown | 7.7 | 12.5 | n/a | 36 | 362.4 | 8231.9 | 1564.5 |

**Table S3.** qPCR standard DNA template information for all five gene targets

| **Target gene** | **Primers** | **Primer reference** | **Standard source** | **Amplicon size (bp)** | **Fragment mass (g/mol)*** |
| --- | --- | --- | --- | --- | --- |
| 16S rRNA bacterial | 341F/518R | Muyzer 1993 | *Thermus thermophilus* | 719 | 444231 |
| 16S rRNA archaeal | 771F/957R | Ochsenreiter et al. 2003 | *Ca.* Nitrosotenuis aquarius | 227 | 140151 |
| AOB *amoA* | amoA1F/amoA2R | Rotthuwe et al. 1997 | *Nitrosomonas europaea* | 491 | 303255 |
| AOA *amoA* | crenamoA23F/  616R | Tourna et al. 2008 | *Ca.* Nitrosotenuis aquarius | 629 | 388505 |
| CMX *amoA* | comaAF/comaAR pooled | Pjevac et al. 2017 | Aquarium samples (4 amplicons pooled for template) | 415 | **269750 |

* Fragment molecular masses correspond to the sequences listed in Table S2 below.

** Calculated based on average molar mass of 650 g/mol/bp as sequence of DNA amplified was variable

**Table S4.** qPCR standard gene sequences with location of target forward and reverse primers indicated in bold.

| Standard source | | Sequence |
| --- | --- | --- |
| *Thermus thermophilus*  341F/518R gene standard | GTAAAACGACGGCCAGTGAATTCGAGCTCGGTACCTCGCGAATGCATCTAGATATCGGATCCCGGGCCCGTCGACTGCAGAGGCCTGCATGCAACGGGCCCCACT**CCTACGGGAGGCAGCAG**TTAGGAATCTTCCGCAATGGGCGCAAGCCTGACGGAGCGACGCCGCTTGGAGGAAGAAGCCCTTCGGGGTGTAAACTCCTGAACCCGGGACGAAACCCCCGACGAGGGGACTGACGGTACCGGGGTAATAGCGCCGGCCAACTCCGTG**CCAGCAGCCGCGGTAAT**ACGGAGGGCGCGAGCGTTACCCGGATTCACTGGGCGTAAAGGGCGTGTAGGCGGCCTGGGGCGTCCCATGTGAAAGACCACGGCTCAACCGTGGGGGAGCGTGGGATACGCTCAGGCTAGACGGTGGGAGAGGGTGGTGGAATTCCCGGAGTAGCGGTGAAATGCGCAGATACCGGGAGGAACGCCGATGGCGAAGGCAGCCACCTGGTCCACCCGTGACGCTGAGGCGCGAAAGCGTGGGGAGCAAACCGGATTAGATACCCGGGTAGTCCACGCCCTAAACGATGCGCGCTAGGTCTCTGGGTCTCCTGGGGGCCGAAGCTAACGCGTTAAGCGCGCCGCCTGGGGAGTACGGCCGCAAGGCTGAAACTCAAAGGAATTGACGGGGGCCCGCACAAAGCTTGGCGTAATCATGGTCATAGCTGTTTCCTG | |
| *Ca.* Nitrosotenuis aquarius  771F/957R gene standard | **ACGGTGAGGGATGAAAGCT**GGGGGAGCAAACCGGATTAGATACCCGGGTAGTCCCAGCTGTAAACGATGCAGACTCGGTGATGCATTGGCTTGTGGCCAATGCAGTGCCGCAGGGAAGCCGTTAAGTCTGCCGCCTGGGAAGTACGTACGCAAGTATGAAACTTAAAGGAATTGGCGGGGGAGCACCACAAGGGGTGAAGCCTGCGGTT**CAATTGGAGTCAACGCCG** | |
| *Nitrosomonas europaea*  amoA1F/amoA2R gene standard | **CCCCTCTGGAAAGCCTTCTTC**ACCGAATGCGGTAACATCATTGCGATGTACGATACGACCTCTTTTACCTTTAACGTAGAAAAAGGCTGTACAGTAAACTTTTCCAAGATACCACCATACGGTGAACATCAACATTGATACGAACGCAGAGAAGAATGCTGCAATAACTGTGGTATGACCACCAAAGGTACGCAGTGAACCTTGCTCAATATGACGAACATACTCGGGTGTACCTGTACGAACATACAGATGTCCCATGTAATCAGCCATCGACAGCAATGTGCCTTCTACAACGATTGGCAAATGGGTTGGTCCAAAAATCGGCCAGTTACCCGGATAGAACAGCAGACCGAAGAATCCACCTCCAACCAGAGCCGTCACCAGCCAGTTGCGTGTCAGATACAGCGTGAAGTCCAGCATCAGCGCACCCGGAAGCATAATGCCCGGTGTTACGAAGTTGATGGGGTAGTGTG**ACCACCAGTAGAATCCCC** | |
| *Ca.* Nitrosotenuis aquarius  crenamoA23F/616R | **ATGGTCTGGCTTAGACG**ATGTACGCACTACTTGTTCATAGTAGTCGTAGCAGTCAACTCAACCCTGCTTACAATCAACGCAGGAGACTACATCTTCTACACTGACTGGGCATGGACTTCGTATGTCGTGTTCTCAATATCACAGACATTGATGTTGGTGGTAGGTGCAACTTACTATCTGACATTTACCGGAGTTCCAGGAACCGCAACATACTACGCGCTGATTATGACCGTGTATACATGGATCGCAAAAGGCGCATGGTTTGCTCTAGGTTACCCATATGACTTCATTGTTACACCAGTTTGGTTACCATCAGCAATGCTGATTGACTTAGCATACTGGGCTACAAAGAAGAACAAGCACTCACTGATACTATTCGGTGGTGTGTTGTGTGGAATGTCACTGCCATTGTTCAACATGGTAAATCTAATTACCGTGGCTGATCCATTGGAGACTGCTTTCAAATATCCAAGACCAACATTGCCTCCATACATGACTCCAATAGAACCCCAAGTGGGCAAGTTCTATAACAGTCCAGTTGCACTCGGTGCAGGCGCAGGCGCTGTATTATCAGTAACCTTTGCCGCTCTGGGATGTAAGCTGAATACG**TGGACGTACAGATGGATGGC** | |

| **Table S5.** Gene copies for both *amoA* and 16S rRNA gene targeted qPCRs for each sample. Copies are expressed per ng of extracted DNA. |  | Comammox *Nitrospira amoA* | | AOB *amoA* | | AOA *amoA* | | Bacterial  16S rRNA gene | | Thaumarcheotal  16S rRNA gene | |
| --- | --- | --- | --- | --- | --- | --- | --- | --- | --- | --- | --- |
| Sample ID | Type | Gene copies per ng gDNA | SD | Gene copies per ng gDNA | SD | Gene copies per ng gDNA | SD | Gene copies per ng gDNA | SD | Gene copies per ng gDNA | SD |
| FW-F01 | Freshwater | 2679.1 | 104.2 | 2.3 | 0.2 | 48.0 | 3.4 | 357650.7 | 48439.3 | 337.9 | 35.8 |
| FW-F02 | Freshwater | 1260.1 | 113.1 | 23.5 | 1.2 | 4.1 | 0.2 | 300566.0 | 46456.8 | 1.5 | 0.6 |
| FW-F03 | Freshwater | 3328.1 | 130.7 | 3.2 | 1.2 | 73.7 | 1.8 | 259100.2 | 24032.9 | 652.1 | 25.4 |
| FW-F04 | Freshwater | 3597.9 | 244.1 | 18.4 | 4.4 | 202.1 | 2.8 | 292563.9 | 41634.0 | 250.3 | 16.5 |
| FW-F05 | Freshwater | 1192.6 | 13.9 | 0.8 | 0.1 | 78.2 | 1.8 | 150118.1 | 5238.3 | 356.4 | 16.9 |
| FW-F06 | Freshwater | 6682.5 | 794.2 | 5.5 | 0.6 | 1062.7 | 225.2 | 277426.1 | 12601.3 | 830.7 | 0.1 |
| FW-F07 | Freshwater | 1396.3 | 84.2 | 46.3 | 8.4 | 2.6 | 1.0 | 582230.8 | 97091.0 | 26.5 | 12.8 |
| FW-F08 | Freshwater | 2351.5 | 4.1 | 1.7 | 0.4 | 1.6 | 0.7 | 378127.9 | 37235.2 | 10.6 | 5.3 |
| FW-F09 | Freshwater | 1730.8 | 223.3 | 6.1 | 2.7 | 65.8 | 12.2 | 375128.7 | 15513.8 | 816.5 | 125.5 |
| FW-F10 | Freshwater | 596.2 | 28.9 | 0.4 | 0.4 | 28.3 | 9.2 | 436530.5 | 69169.3 | 30.5 | 12.0 |
| FW-F11 | Freshwater | 1235.2 | 159.7 | 0.9 | 0.4 | 0.7 | 0.1 | 180345.5 | 2670.1 | 42.3 | 5.6 |
| FW-F12 | Freshwater | 2453.8 | 98.4 | 1.4 | 0.1 | 13453.9 | 1552.3 | 381424.0 | 81179.0 | 21162.7 | 4009.6 |
| FW-F13 | Freshwater | 3761.0 | 2.4 | 12.3 | 3.3 | 17.1 | 0.8 | 238815.4 | 1040.4 | 99.7 | 14.1 |
| FW-F14 | Freshwater | 5010.0 | 1046.4 | 1.7 | 0.2 | 2368.9 | 210.1 | 240915.5 | 44783.9 | 1792.9 | 210.7 |
| FW-F15 | Freshwater | 1455.3 | 131.4 | 0.5 | 0.1 | 1976.5 | 279.9 | 334006.2 | 267705.7 | 2488.2 | 370.2 |
| FW-F16 | Freshwater | 1445.2 | 216.3 | 1.6 | 0.1 | 15.6 | 1.6 | 354042.2 | 8000.5 | 146.9 | 9.5 |
| FW-F17 | Freshwater | 2364.1 | 447.8 | 2.2 | 0.2 | 256.3 | 33.6 | 612831.7 | 136570.8 | 405.7 | 102.7 |
| FW-F18 | Freshwater | 842.5 | 61.6 | 0.5 | 0.1 | 1.8 | 0.1 | 106739.5 | 19987.6 | 2.0 | 0.2 |
| FW-F19 | Freshwater | 645.3 | 83.0 | 0.6 | 0.2 | 70.4 | 12.9 | 238073.3 | 70781.2 | 356.5 | 95.3 |
| FW-F20 | Freshwater | 960.3 | 58.1 | 109.0 | 10.4 | BDL | - | 100728.4 | 3753.2 | 0.5 | 0.0 |
| FW-F21 | Freshwater | 545.3 | 21.1 | 1.7 | 0.1 | 358.6 | 11.3 | 386154.4 | 109426.1 | 524.1 | 16.6 |
| FW-F22 | Freshwater | 1179.2 | 130.8 | 138.1 | 31.9 | 8.8 | 2.3 | 602839.7 | 28901.7 | 474.5 | 31.6 |
| FW-F23 | Freshwater | 4368.9 | 77.4 | 689.0 | 196.6 | 116.7 | 6.4 | 1210216.3 | 237183.9 | 202.6 | 36.6 |
| FW-F24 | Freshwater | 1765.9 | 227.4 | 12.1 | 2.2 | 2019.4 | 223.8 | 1307997.8 | 1389547.8 | 2093.8 | 172.0 |
| FW-F25 | Freshwater | 461.4 | 36.7 | 2.1 | 0.4 | 646.3 | 9.3 | 101620.5 | 23826.8 | 1310.2 | 178.9 |
| FW-F26 | Freshwater | 2238.6 | 476.5 | 8.3 | 0.2 | 165.9 | 32.3 | 566035.2 | 153888.7 | 1056.8 | 32.0 |
| FW-F27 | Freshwater | 2786.2 | 430.2 | 3.5 | 0.0 | 178.4 | 46.7 | 463383.5 | 30854.2 | 437.5 | 88.3 |
| FW-F28 | Freshwater | 884.1 | 130.8 | 2.0 | 0.3 | 2359.7 | 357.4 | 149997.0 | 7441.8 | 1968.6 | 584.8 |
| FW-F29 | Freshwater | 4973.7 | 957.6 | 4.9 | 0.6 | 1134.3 | 217.1 | 545152.5 | 51591.8 | 1805.4 | 263.4 |
| FW-F30 | Freshwater | 2926.2 | 398.5 | 1.4 | 0.2 | 462.2 | 4.9 | 267397.7 | 35865.0 | 512.1 | 55.5 |
| FW-F31 | Freshwater | 1674.7 | 118.6 | 2.5 | 0.6 | 2575.8 | 343.2 | 121920.8 | 14029.9 | 743.9 | 65.6 |
| FW-F32 | Freshwater | 1570.9 | 0.5 | 1.4 | 0.1 | 138.5 | 3.6 | 311943.9 | 22899.5 | 264.6 | 35.4 |
| FW-F33 | Freshwater | 1138.0 | 112.3 | 5.9 | 0.1 | BDL | - | 387222.8 | 149070.7 | 2.5 | 0.3 |
| FW-F34 | Freshwater | 34.6 | 0.0 | 68.3 | 10.4 | BDL | - | 176799.5 | 7174.0 | 2.2 | 0.2 |
| FW-F35 | Freshwater | 1467.0 | 301.3 | 4.5 | 0.4 | 1.2 | 0.0 | 957348.9 | 799163.3 | 10.3 | 4.4 |
| FW-F36 | Freshwater | 4729.0 | 383.1 | 3.9 | 0.9 | 9922.1 | 542.0 | 432699.9 | 68186.8 | 14471.5 | 2056.0 |
| FW-F37 | Freshwater | 3233.5 | 296.8 | 15.3 | 1.4 | 779.2 | 62.0 | 119497.4 | 14233.0 | 338.1 | 4.8 |
| FW-F38 | Freshwater | 1594.3 | 179.4 | 0.8 | 0.5 | 2.1 | 0.9 | 136777.2 | 23155.9 | 40.0 | 4.9 |
| SW-F01 | Saltwater | BDL | - | 218.5 | 8.4 | 631.1 | 81.2 | 204763.7 | 45638.6 | 4830.5 | 364.3 |
| SW-F02 | Saltwater | BDL | - | 348.4 | 14.2 | 354.3 | 42.3 | 166481.8 | 36453.6 | 6319.1 | 1725.6 |
| SW-F03 | Saltwater | BDL | - | 2.3 | 0.9 | 69.0 | 16.3 | 137252.7 | 29006.7 | 2367.6 | 57.8 |
| SW-F04 | Saltwater | BDL | - | 107.1 | 6.4 | 194.3 | 15.2 | 240120.1 | 14018.5 | 3476.7 | 411.5 |
| SW-F05 | Saltwater | BDL | - | 261.3 | 12.8 | 29.9 | 2.2 | 393031.5 | 238249.5 | 896.2 | 233.9 |
| SW-F06 | Saltwater | BDL | - | 264.3 | 29.2 | 59.2 | 2.4 | 286943.4 | 18283.7 | 165.9 | 12.5 |
| SW-F07 | Saltwater | BDL | - | 393.7 | 96.9 | 596.8 | 8.5 | 240783.8 | 17034.1 | 8470.4 | 516.0 |
| SW-F08 | Saltwater | BDL | - | 18.5 | 3.3 | 28.4 | 5.8 | 263155.1 | 4273.9 | 480.5 | 69.6 |
